# A Medium Chain Fatty Acid, 6-hydroxyhexanoic acid (6-HHA), Protects Against Obesity and Insulin Resistance

**DOI:** 10.1101/2023.07.19.549684

**Authors:** Sara C. Sebag, Qingwen Qian, Chawin Upara, Qiong Ding, Huojun Cao, Liu Hong, Ling Yang

**Author notes:** Address all correspondence to: Ling Yang, Ph.D. Department of Anatomy and Cell Biology Fraternal Order of Eagles Diabetes Research Center Pappajohn Biomedical Institute University of Iowa Carver College of Medicine, Iowa City, IA 52242 Or: Liu Hong, Ph.D., MD. Iowa Institute for Oral Health Research Department of Prosthodontics, University of Iowa College of Dentistry, Iowa City, 52242.

## Abstract

Obesity, a worldwide health problem, increases the risk for developing metabolic diseases such as insulin resistance and diabetes. It is well recognized that obesity-associated chronic inflammation plays a key role in the pathogenesis of systemic metabolic dysfunction. Previously, we revealed an anti-inflammatory role for spent culture supernatants isolated from the oral commensal bacterial species *Streptococcus gordonii (Sg-SCS).* Here, we identified that 6-hydroxyhexanoic acid (6-HHA), a medium chain fatty acid (MCFA), is the one of the key components of *Sg-SCS*. We found that treatment of 6-HHA in mice fed a high-fat diet (HFD) significantly reduced HFD-mediated weight gain which was largely attributed to a decrease in fat mass. Systemically, 6-HHA improves obesity-associated glucose intolerance and insulin resistance. Furthermore, administration of 6-HHA suppressed obesity-associated systemic inflammation and dyslipidemia. At the cellular level, treatment of 6-HHA ameliorated aberrant inflammatory and metabolic transcriptomic signatures in white adipose tissue of mice with diet-induced obesity (HFD). Mechanistically, we found that 6-HHA suppressed adipocyte-proinflammatory cytokine production and lipolysis, the latter through Gαi-mediated signaling. This work provides direct evidence for the anti-obesity effects of a novel MCFA, which could be a new therapeutic treatment for combating obesity.

**KEY POINTS:**

- Hydroxyhexanoic medium chain fatty acids (MCFAs) are dietary and bacterial-derived energy sources, however, the outcomes of using MCFAs in treating metabolic disorders are diverse and complex.
- The MCFA 6-hydroxyhexanoic acid (6-HHA) is a metabolite secreted by the oral bacterial commensal species *Streptococcus gordonii;* here we investigated its role in modulating high-fat diet (HFD)-induced metabolic dysfunction.
- In a murine model of obesity, we found 6-HHA-mediated improvement of diet-mediated adiposity, insulin resistance and inflammation were in part due to actions on white adipose tissue (WAT).
- 6-HHA suppressed proinflammatory cytokine production and lipolysis through Gi-mediated signaling in differentiated white adipocytes.

## INTRODUCTION

Obesity, one of the most serious health concerns worldwide, is a risk factor for several debilitating illnesses including diabetes and cardiovascular disease (1). Even though lifestyle changes, including exercise and caloric restriction, have been shown to be effective for short-term weight loss, long-term body weight control/management remains a challenge, thus obesity persists/recurs due to weight regain. Therefore, alternative strategies are needed for sustainable weight management.

Although obesity is characterized by dyslipidemia, certain dietary lipids are associated with beneficial outcomes for diabetic and obese patients (2-7). For instance, supplementation of dietary polyunsaturated fatty acids (PUFAs) results in anti-obesogenic and -atherosclerotic effects in humans (8, 9). Furthermore, branched fatty acyl esters of hydroxyl fatty acids (FAHFAs, including short-, medium- and long-chain fatty acids), have been shown to contain antidiabetic properties (10, 11). Finally, coconut oil and dairy products-derived hydroxyhexanoic medium chain fatty acids (MCFAs) are important substrates and/or intermediates of energy metabolism (11). Characterized by a chain length of 6-12 carbons, MCFAs are absorbed predominantly via the portal vein into the liver bypassing the lymphatic system (12), and MCFAs are water-soluble in the intestinal lumen and cytoplasm of target cells. Physiologically, MCFAs have been shown to modulate immune cell function (13), reduce inflammatory responses (14-16), serve as antioxidant (17), and activate ligand-dependent transcription factors involved in insulin sensitivity in diverse set of cell types (18, 19). Further, in humans and rodents, *in vivo* studies have implicated potential role of MCFAs in positively regulating oxidative metabolism and reducing adiposity (2, 20-23). Thus, there is a need/significance to understand the complex regulatory roles, functions, and implications of FA supplementation on human health and disease.

In addition to dietary lipid supply, the oral and gut microbiota produce FAs as end-products of their metabolism. These metabolites function to promote “cross-feeding” between bacterial species and alter microbial ecology (24, 25), provide the host with essential nutrients and aid in defense against colonization of opportunistic pathogens through modulation of the host’s immune system (26-29). It is well recognized that gut and oral-derived short chain fatty acid end-products (SCFA) play a role in health and obesity (27, 30-34); however, the function of MCFAs is largely unknown. Recently, we demonstrated that secretions from the oral commensal bacterial species *Streptococcus gordonii* protect against pathogen-mediated inflammation of oral epithelial cells (35, 36). In this study we identified that one of these secreted components is a medium chain, omega-hydroxy fatty acid, 6-hydroxyhexanoic acid (6-HHA). Here we investigated the pathophysiological relevance of 6-HHA in the context of obesity. We found that 6-HHA improves inflammation and insulin resistance in mice with diet-induced obesity (DIO) while decreasing FFA release and pro-inflammatory mediators from white adipocytes. These findings demonstrate the potential for novel FAs in management of diet-induced obesity and associated comorbidities.

## METHODS

### Cell Culture and treatment

3T3-L1 (CL-173-ATCC, (37)) were grown in DMEM containing 10% fetal bovine serum (FBS) and 1% pen/strep. Cells were induced to differentiate 2 days after reaching confluence by supplementing growth media with 3nM insulin (Humulin R, Lily, Humulin R U-500, 0002-8501-01), 0.25nM dexamethasone (Sigma, D4902), 2uM rosiglitazone (Sigma, R2408) and 0.5mM 1-methyl-3-isobutyl-xanthine (Sigma, I5879). From day 3 until day 7, cells were maintained in growth media supplemented with 3nM insulin after which the mature adipocytes were maintained in growth media. Cells were then incubated for 24 hr in the presence or absence of 10ng/ml TNF (PeproTech, 315-01A) with or without 0.1uM 6-Hydroxyhexanoic acid (6-HHA, ThermoFisher, B24857.03).

For Free Fatty Acid (FFA) measurement, cells were incubated with phenol-free media containing 5% FFA-BSA. Cells were pretreated with 50ng/ml Pertussis Toxin (Enzo, BML-G101-0050) 24 hr prior to 0.1uM 6-HHA or media alone (24 hr). Supernatants were collected, fresh media was added and then collected again at 20 min. Finally, cells were treated with 1uM isoproterenol (Sigma, I6504) for 1 hr. Supernatant and cell lysates were collected and frozen until assessed.

### Animal Experiments

Animal care and experimental procedures were performed with approval from the University of Iowa’s Institutional Animal Care and Use Committee. Animals received humane care in compliance with the *Guide for the Care and Use of Laboratory Animals* (National Academies Press, 2011) and with the Principles of Laboratory Animal Care formulated by the National Society for Medical Research.

C57BL/6J mice (The Jackson Laboratory, 000664) were kept on a 12-hour light/dark cycle. Mice used to generate the DIO model were placed on a 60% kCal high-fat diet (HFD, Research Diets, D12492) immediately after weaning (i.e., at 3 weeks of age). After 16 weeks, lean and DIO-mice were intraperitoneally (IP) injected with 0.25 or 0.5uM 6-HHA or PBS control every other day for three weeks. Body weight was measured weekly.

All tissues were harvested, frozen in liquid nitrogen, and kept at −80°C until processed.

### Quantitative Real-time RT-PCR

Total RNA was isolated using TRIzol reagent (Invitrogen,15-596-018) and reverse transcribed into cDNA using the iScript cDNA synthesis kit (Bio-Rad, 1708890). Quantitative real-time RT-PCR analysis was performed using SYBR Green (Invitrogen, KCQS00). Mouse Gapdh: Forward: TGTGTCCGTCGTGATCTGA; Reverse: CCTGCTTCACCACCTTCTTGAT. Mouse Nos2: Forward: GTTCTCAGCCCAACAATACAAGA; Reverse: GTGGACGGGTCGATGTCAC. Mouse Adipoq: Forward: GCACTGGCAAGTTCTACTGCAA; Reverse: GTAGGTGAAGAGAACGGCCTTGT. Mouse PParg Forward: TATGGAGTGACATAGAGTGTGCT; Reverse: CCACTTCAATCCACCCAGAAAG. Mouse Lep: Forward: GAGACCCCTGTGTCGGTTC; Reverse: CTGCGTGTGTGAAATGTCAATG. Mouse Srebp1c Forward: CTGCATAACGGTCTGGACTTC; Reverse: GGCCCGGGAAGTCACTGT. Mouse FasN: Forward: AGAGATCCCGAGACGCTTCT; Reverse: GCCTGGTAGGCATTCTGTAGT. Mouse Il1b: Forward: GCAACTGTTCCTGAACTCAACT; Reverse: ATCTTTTGGGGTCCGTCAACT. Mouse Gpr81: Forward: GGTGGCACGATGTCATGTT; Reverse: GACCGAGCAGAACAAGATGATT. Mouse Cd36: Forward: GAGCCATCTTTGAGCCTTCT. Reverse: TCAGATCCGAACACAGCGTA. Mouse Hsl: Forward: AGCCTCCCACATGCGTTCTGC. Reverse: GCGAGGTGTCTCTCTGCACGA. Mouse Il6: Forward: CTGCAAGAGACTTCCATCCAG. Reverse: AGTGGTATAGACAGGTCTGTTGG. Mouse Atgl: Forward: CCAGCATCCAGTTCAACCTTC. Reverse: GGCTTAGTCAGGACTGGGTGT.

### RNA-seq and data analysis

For each test group, total RNA was extracted from frozen WAT samples from 3 mice. Generation of the RNA-seq library and sequencing were performed by the University of Iowa Genomics Core. RNA-seq reads were quality checked using the FastQC tool. Low-quality and adapter sequences were trimmed using the Trimmomatic (38). Expression of transcripts was quantified using the Salmon tool (39) and estimates of transcript abundance for gene-level analysis were imported and summarized using the tximport (40) function of the R/Bioconductor software suite (41). Differentially expressed genes (DEGs) were identified by applying the R/Bioconductor package Deseq2 (42). Enriched pathways represented by the DEGs were identified by gene set enrichment analysis (GSEA) (43) and Enrichr (44).

### Western blot Analysis

Proteins were extracted from cells or tissues and subjected to SDS–polyacrylamide gel electrophoresis, as previously described (45). Membranes were incubated with anti-p-AKT (Ser473, Cell Signaling, 9271); anti-AKT1 (H-136, Santa Cruz, sc-8312) or anti-ACTB (H-300, Santa Cruz, sc-10731) at 1:1000 and then incubated with the appropriate secondary antibody conjugated with horseradish peroxidase (1:5000, Santa Cruz, sc-2005 or 1:5000, Cell Signaling Technology, 7074S). Signal was detected using the ChemiDoc Touch Imaging System (Bio-Rad), and densitometric analyses of western blot images were performed using Image Lab software (Bio-Rad).

### Immunohistochemistry

For immunohistochemistry, tissues were fixed with 4% PFA and sectioned at 5 μm thick, followed by deparaffinization and rehydration processes. Tissue sections were stained using H&E. The images were observed under a Nikon microscope (10x).

### Biochemical analysis and cytokine measurements

Blood samples were centrifuged (4°C, 5000*g*, 30 min) to obtain serum. Ten microliters of serum per sample were required for each index analysis. Serum alanine aminotransferase (ALT) and aspartate aminotransferase (AST) were measured by commercial kits. Serum IL-1*β*, IL-6 and FFA content were measured by ELISA (IL-1*β*, Biolegend, 432604; IL-6, Invitrogen, BMS603-2) or a FFA fluorometric kit (Cayman Chemical, 700310), respectively, and then normalized to total protein (BCA).

### Metabolic Phenotyping

*Whole-body energy expenditure and body composition*: Respiratory exchange ratio (RER) and locomotor activity were monitored using a Comprehensive Lab Animal Monitoring System (CLAMS, Columbus Instruments) at the Fraternal Order of Eagles Metabolic Phenotypic Core. Body composition was measured using Bruker Minispecs (LF50).

### Insulin tolerance test (ITT)

Animals were fasted for 6 hours prior to ITT. Insulin tolerance was tested by measuring glucose concentration at different time points after intraperitoneal (IP) insulin injection (0.5U/kg body weight; Humulin R) (46).

### Statistical Analysis

Results are expressed as the mean ± the standard error of the mean (SEM); *n* represents the number of individual mice (biological replicates) or individual experiments (technical replicates) as indicated in the figure legends. We performed the Shapiro-Wilk Normality test in experiments that have a relatively large sample size (n>5) and found that these data pass the normality test (alpha=0.05). Data were further analyzed with two-tailed Student’s and Welch’s *t*-test for two-group comparisons or ANOVA for multiple comparisons. For both One-Way ANOVA and Two-Way ANOVA, Tukey’s post-hoc multiple comparisons were applied as recommended by Prism. In all cases, GraphPad Prism (GraphPad Software Prism 8) was used for the calculations.

## RESULTS

### 6-hydroxyhexanoic acid (6-HHA) ameliorates obesity-associated metabolic dysfunction

We have previously shown that the spent culture supernatants (SCS) of *Streptococcus gordonii* mitigates inflammation of human periodontal cells and inhibits proliferation of pathogenic oral microbes (35). Further investigation into the components comprising the SCS of *S.gordonii* revealed the presence of a significant level of the metabolite, 6-hydroxyhexanoic acid (6-HHA, shown as HHA in figures), a medium chain fatty acid (MCFA) by high performance liquid chromatography (HPLC) analysis (data not shown). To answer whether and to what extent 6-HHA affect obesity-associated metabolic defects, wildtype (WT) mice fed a high-fat diet (HFD) were injected intraperitoneally (IP) with 6-HHA for three weeks. Compared to mice on RD, mice fed a HFD for 16 weeks significantly gained more body weight (BW) (Fig. 1A). Dose-dependent studies demonstrated HFD-fed mice exposed to 0.25 µM 6-HHA had comparable effects compared to HFD animals (data not shown). However, BW gain was significantly reduced in HFD-fed mice treated with 0.5 µM 6-HHA compared to the vehicle treated HFD mice (Fig. 1A&B), therefore, experiments were performed with 0.5uM 6-HHA. Body composition analysis further revealed that the 6-HHA-mediated reduction in BW gain was mainly due to a decrease in fat mass and slight increases in lean mass (Fig. 1C).

**Figure 1.**
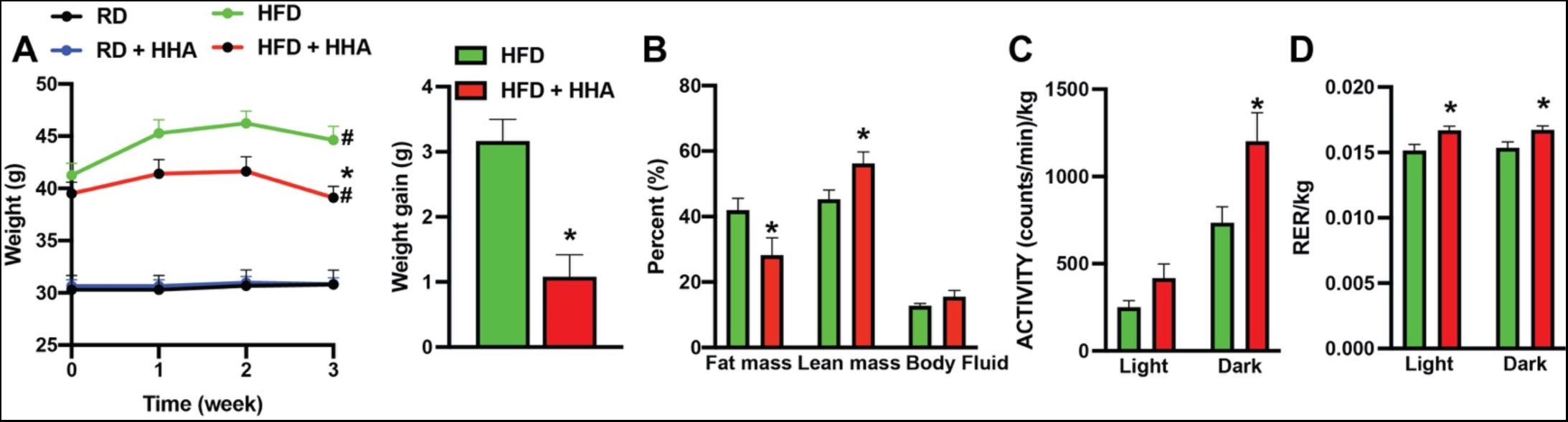
6-HHA protects against obesity-associated metabolic dysfunction. Mice on a regular (RD) or high-fat diet (HFD,16 weeks) were intraperitoneally (IP) injected with control (water) or 6-HHA (0.5 µM) every other day for three weeks. **(A)** Body weight**, (B)** Body composition measured by NMR, **(C)** Total activity and **(D)** respiratory exchange ratio (RER) measured in mice housed within a CLAMS. N = 4-12 mice/group. All data are presented as means ± SEM. *indicates statistical significance of treatment effect between HFD groups, and #indicates statistical significance of dietary effect; as determined by Student’s t-test **(C, D)** or ANOVA followed by post hoc test **(A, B)**, p<0.05.

Obesity disrupts systemic energy balance. Therefore, we measured whole-body metabolic profiles in HFD mice in the presence or absence of 6-HHA by comprehensive lab animal monitoring system. 6-HHA treatment increased activity and shifted the respiratory exchange ratio (RER) towards increased carbohydrate metabolism in HFD mice, suggesting better fuel utilization as measured by indirect mouse calorimetry (Fig. 1C&D). HFD-mediated obesity is associated with impaired systemic glucose homeostasis (47). We next determined whether 6-HHA modulates obesity-associated hyperglycemia and insulin resistance. As shown in Fig. 2A, compared to RD mice, fasting glucose in HFD-fed animals was significantly increased; furthermore, insulin tolerance was impaired (Fig. 2A&B). While 6-HHA did not alter glucose and insulin homeostasis in RD mice, it significantly improved obesity-associated hyperglycemia and insulin resistance, without altering serum insulin levels (Fig. 2A-C). Together, these data demonstrated that 6-HHA protects against obesity-associated systemic energy and glucose imbalance.

**Figure 2.**
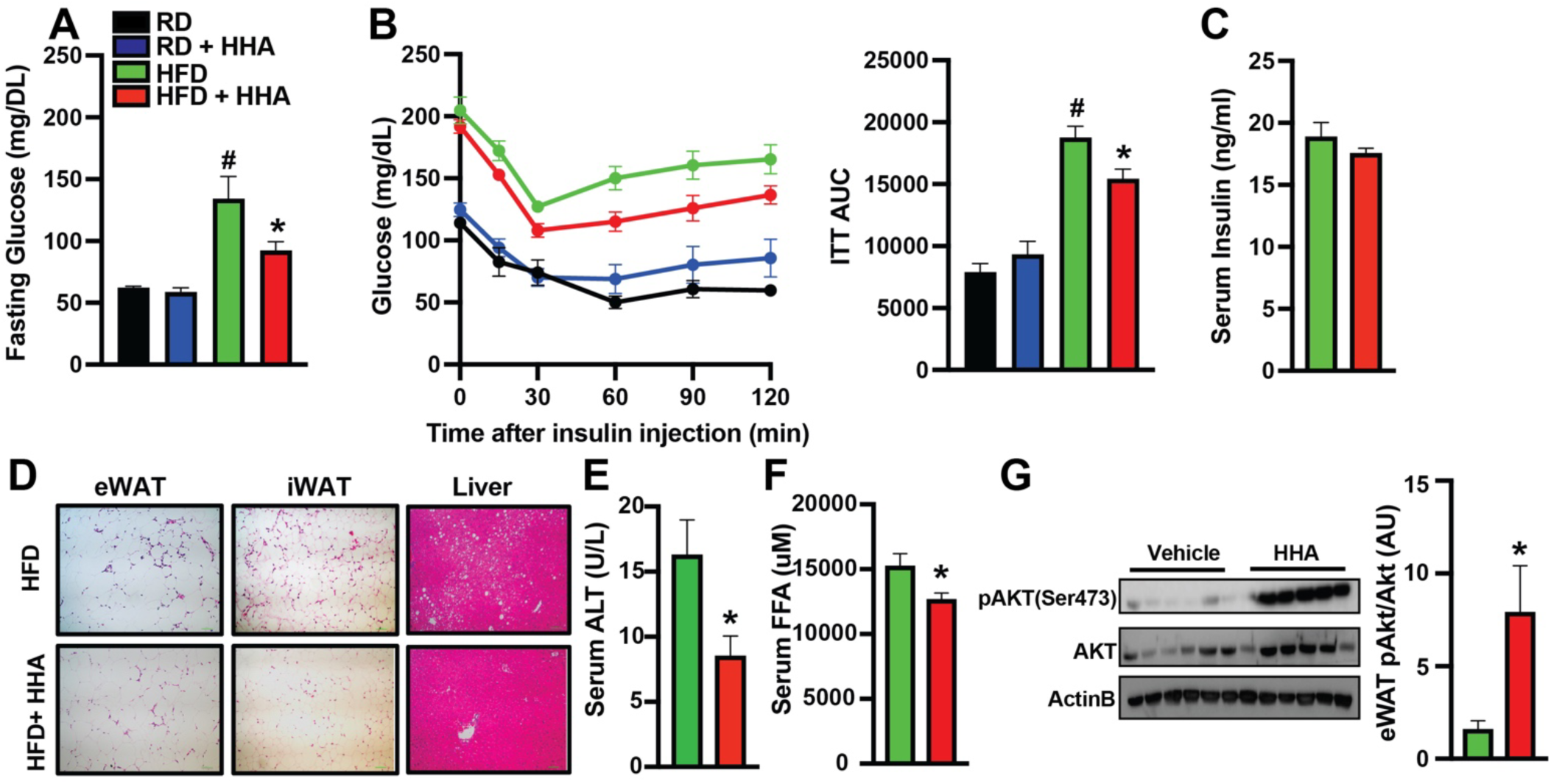
6-HHA improves systemic glucose homeostasis and insulin sensitivity in obese mice. Mice were fed RD or HFD followed by IP injection with control (water) or 6-HHA (0.5 µM) every other day for three weeks. **(A)** 16-hr fasting glucose levels, **(B)** Insulin tolerance test (ITT), **(C)** Serum insulin levels, and **(D)** Representative light microscopy and H&E images (20X) in eWAT, inguinal white adipose (iWAT) and the livers of the experimental mice. Scale bar: 10μm. (**E)** Serum alanine transaminase (ALT) or **(F)** free fatty acid (FFA) were measured in mice from **(A).** N = 6 mice/group. **(G)** Insulin signaling in white adipose tissue of mice as in **(A)** p-AKT: AKT^ser473^. Densitometric analysis is on the right panel, and data are normalized to ACTB. N = 4-12mice/group. All data are presented as means ± SEM. *indicates statistical significance of treatment effect between HFD groups, and #indicates statistical significance of dietary effect; as determined by one-way ANOVA **(A, and B** of AUC) and by Student’s t-test **(E, F and G),** p<0.05.

### 6-HHA improves metabolic tissue function in obese mice

In obesity, the adipose tissues and liver become dysfunctional, resulting in impaired glucose and lipid metabolism (48, 49). To evaluate the actions of 6-HHA at the tissue/organ level, we first assessed hepatic and adipose lipid accumulation in the liver and white adipose depots from HFD and 6-HHA-treated HFD mice. We found that 6-HHA treatment lowered intra-organ fat deposit in the liver, inguinal and epididymal white adipose tissue (iWAT and eWAT, respectively) (Fig. 2D). Obesity-induced hepatic steatosis leads to liver damage (49). To assess liver toxicity associated with 6-HHA, we measured serum alanine transaminase (ALT). Consistent with the decreased hepatic steatosis, we show that 6-HHA treatment improved obesity-associated liver damage (Fig. 2E). Additionally, liver transcripts involved in lipogenesis, lipolysis and inflammation were altered in 6-HHA treated mice (SFig. 1A). Elevated circulating levels of free fatty acids (FFAs), caused by increased basal lipolysis in adipose tissues, are a primary reason for the development of insulin resistance in metabolic disorders (50). We then measured serum level of free fatty acid (FFA) in HFD and HFD with or without 6-HHA treatment and found 6-HHA decreased lipolysis compared to HFD controls (Fig. 2F). These data indicate 6-HHA modifies hepatic lipolytic activity in DIO mice. Adipose insulin resistance is the major driver for obesity-associated lipolysis (51). Indeed, we found that 6-HHA treatment significantly improved adipose insulin sensitivity in eWAT of HFD-fed mice (Fig. 2G).

### The impact of 6-HHA on adipose transcriptome in the context of obesity

To investigate the HFD-mediated genome-wide changes in RNA levels in the eWAT mediated by 6-HHA, we performed RNA-seq. Signaling pathways involved in fatty acid metabolism, PPARgamma as well as AMPK were significantly increased while chemokine signaling and leukocyte migration were downregulated (Fig. 3A&B). Additionally, HFD-mediated fatty acid metabolism and biosynthesis were significantly modified by 6-HHA exposure (Fig. 3C&D).

**Figure 3.**
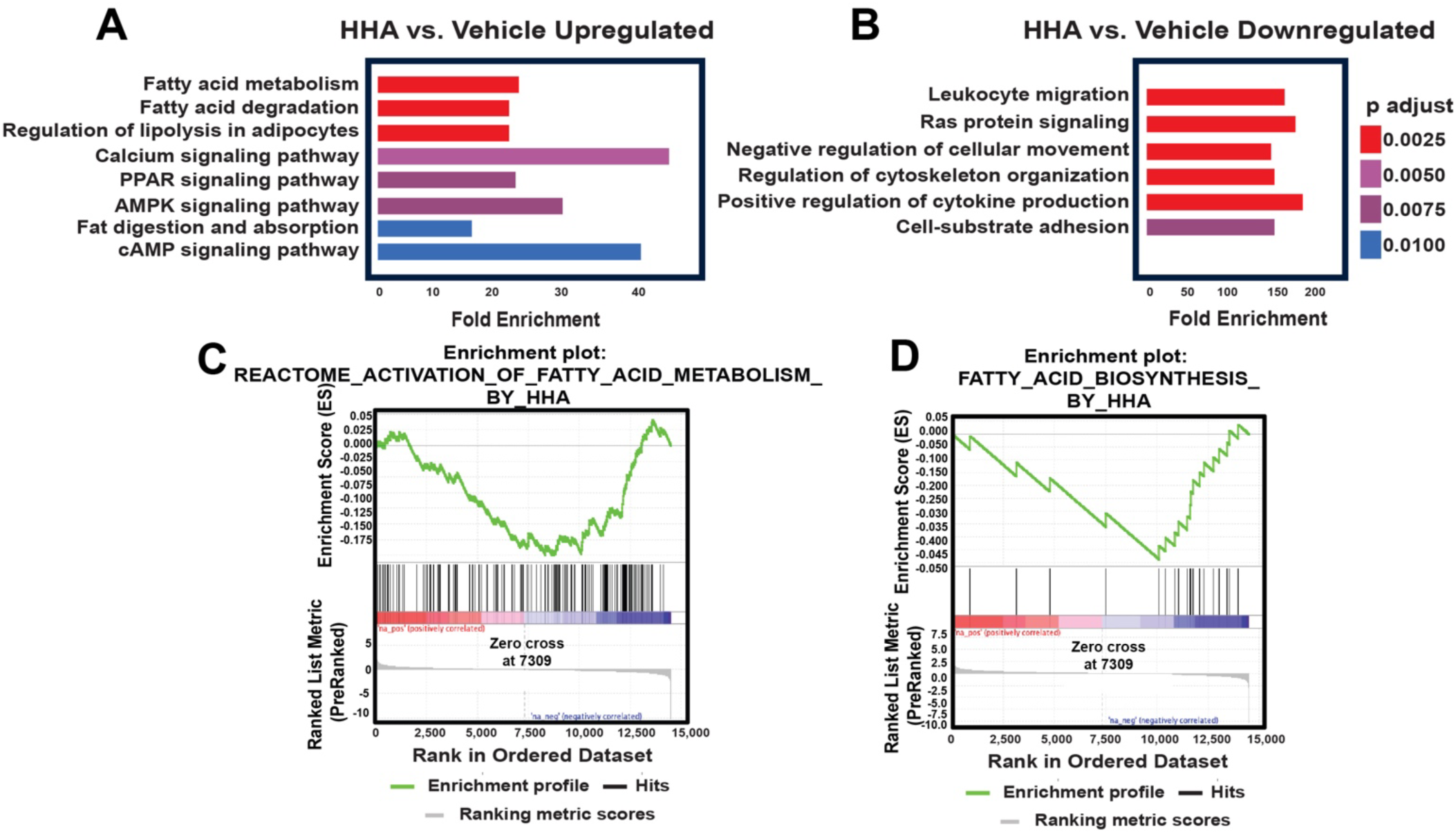
6-HHA alters white adipose tissue metabolic and inflammatory signature. RNA-seq from white adipose tissue from mice fed HFD followed by IP injection with control (water) or 6-HHA (0.5 µM) every other day for three weeks. Pathways that are significantly **(A)** up- or **(B)** downregulated; KEGG categories are determined by Enrichr analysis. Gene Set Enrichment Analysis (GSEA) plots illustrate significant activation of fatty acid metabolism by 6-HHA **(C)** or fatty acid biosynthesis **(D).** N= 4 mice/group.

### 6-HHA improves obesity-associated adipose inflammation and metabolic defects

To establish the cellular mechanism underlying the protective role of 6-HAA in the adipose tissue in the context of obesity, we next analyzed the expression of several genes involving adipose inflammation and function in eWAT from RD and HFD mice with or without 6-HHA treatment. As shown in Fig. 4A, obesity-induced pro-inflammatory (*Il6*, *Nos2*) as well as white adipose transcript markers (*Lep*) were significantly reduced, while *Adipoq* and *Ppargamma* levels were increased by 6-HHA. Notably *Cd36* and *Gpr81*, receptors involved in FA uptake (52, 53), were significantly elevated by 6-HHA in HFD (Fig. 4A).

**Figure 4.**
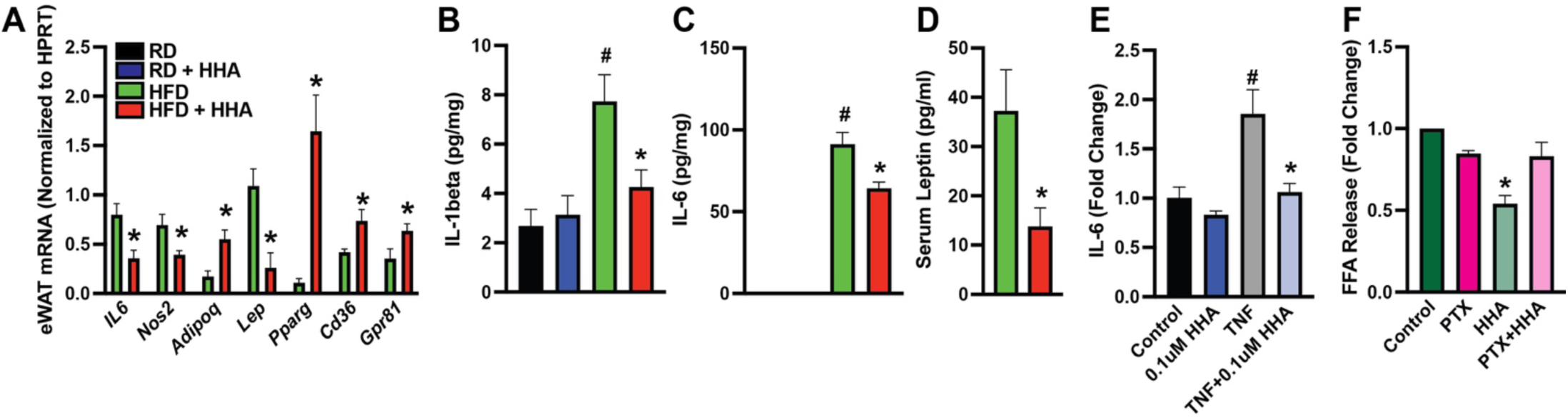
6-HHA improves obesity-associated adipose inflammation and lipolysis. Mice on a RD or HFD (16 weeks) were IP injected with control (water) or 6-HHA (0.5 µM) every other day for three weeks. **(A)** Levels of mRNAs encoding the indicated genes in WAT, data were normalized to *Gapdh.* Serum levels of **(B)** IL-1beta, **(C)** IL-6 and **(D)** leptin in mice as in **(A).** N = 4-6 mice/group. **(E)** IL-6 production in differentiated 3T3L-1 cells treated with in 0.1 µM 6-HHA (24 hr) with in the presence or absence of 10ng/ml TNF. Values were normalized to total protein (BCA). **(F)** FFA releasing from differentiated 3T3L-1 cells, values were normalized to total protein (BCA). N= 3 (experimental replicates). All data are presented as means ± SEM. *indicates statistical significance of treatment effects, and ^#^indicates statistical significance of dietary effect; as determined by Student’s t-test **(A and D),** one-way ANOVA **(B, C, E, F)**, p<0.05.

Obesity-promoted adipose inflammation is the major driver for obesity-associated metabolic dysfunction (54). We then measured serum levels of inflammatory mediators often associated with obesity and insulin resistance. Notably, the 6-HHA-supressed proinflammatory gene expressions were concomitant with reduced serum levels of IL-6, IL-1*β* and leptin (Fig. 4B-D).

Finally, to address the cell autonomous effect of 6-HHA on adipocyte-derived cytokine production, we utilized an adipocyte cell line, 3T3-L1. 3T3-L1 cells were differentiated into white adipocytes and then exposed to the pro-inflammatory cytokine tumor necrosis factor alpha (TNF-α). Indeed, we found TNF-induced IL-6 secretion was significantly reduced by 6-HHA (Fig. 4E).

Adipose-tissue localized G-protein coupled receptors (GPCRs) play important roles in modulation of lipolysis (55), therefore, we then asked whether the inhibitory role of 6-HHA on lipolysis acts through GPCR signaling. Differentiated 3T3-L1 cells were pretreated with pertussis toxin (PTX), which inhibits subunits from the Gα_i_ family (56), prior to 6-HHA exposure. We found that isoproterenol (ISO)-induced FFA release was significantly reduced by 6-HHA in mature adipocytes, however, cotreatment with PTX abrogated this effect (Fig. 4F), indicating a role for Gα_i-_mediated signaling in ISO-mediated adipocyte-FFA release.

## DISCUSSION

Obesity exacerbates periodontitis by increasing its prevalence, accelerating its progression and reducing the effectiveness of treatments (57-59). In turn, periodontitis increases serum lipid and glucose levels, resulting in increased risk of developing insulin resistance, thereby worsening the effects of obesity (60, 61). We have previously shown that oral *S. gordonii*-derived metabolites protect against pathogen-mediated inflammation of oral epithelial cells (35). Here, we demonstrated the protective role of one of the top-enriched components, 6-hydroxyhexanoic acid (6-HHA), on murine model of diet-induced obesity.

Previous studies in mice have shown dietary MCFAs improve DIO-mediated insulin sensitivity (62-64), promote weight loss (65, 66) and inhibit inflammatory responses (47). Here, we found that 6-HHA, an oral microbiota-derived MCFA, ameliorates adiposity and systemic inflammation and improves systemic energy and metabolic homeostasis in DIO mice. We further revealed the direct regulatory role of 6-HHA on adipose metabolic function. However, the effects of MCFAs on glucose homeostasis in human patients remain contradictory. For instance, in lean men, dietary intake of MCFA increased serum insulin while reducing glucose levels (67) or had no effect (68). In conditions of overnutrition, MCFAs rescued insulin action in lean humans (69) whereas in patients with non-insulin-dependent diabetes mellitus (NIDDM), dietary intake of MCFA had either no effect or improved insulin-mediated glucose clearance (68, 70, 71).

Regarding the interplay between the oral and gut microbiota and obesity, a large body of studies have primarily focused on modulation of microbial species in the context of metabolic syndrome (72-74). It has been shown that microbe-derived branched-chain fatty acids are implicated in the pathogenesis of obesity and diabetes, although findings are controversial and complex (75, 76). For instance, a study demonstrated another commensal species, *Fusimonas intestine (FI)* which produces long-chain fatty acids (LCFA), is increased in the gut of obese human subjects and exacerbates mouse models of DIO (77). However, in other models, gut-mediated microbiota protects against high-fat diet (HFD)-mediated obesity (78, 79). Furthermore, although specific oral-derived bacteria vary with obesity in both adolescents and adults, little is known about their roles in mediating the pathogenesis of obesity (57, 80, 81). Thus, there is a need to better understand the role of FA-produced, oral microbial metabolites on preventing or reducing the prevalence of obesity and associated metabolic disorders.

Pathologically, DIO elevates plasma lipids which in turn increase FA accumulation in non-adipose tissues resulting in insulin resistance (82). We found 6-HHA treatment resulted in reduced fat mass, increased nocturnal activity and altered fuel utilization. Notably, to further understand the role of 6-HHA on obesity-mediated dyslipidemia, research should focus on 6-HHA-mediated effects on leptin resistance and metabolic tissue-mediated intracellular lipid utilization. Additionally, DIO-mediated inflammation leads to lipid disorders, obesity and systemic glucose/insulin impairment (54, 83, 84). Our study demonstrated an anti-inflammatory role of 6-HHA both systemically and in adipose tissue in the context of obesity. We recognize this effect might be attributed to the action of 6-HHA on modulating function of immune cells, which is an area of interest for future studies.

In summary, we have revealed a novel oral-derived bacterial commensal metabolite, 6-HHA, plays an essential role in protecting against obesity-associated metabolism dysfunctions. Therefore, supplementing or replacing diets with MCFAs may result in lower body fat mass, improved adipose tissue-mediated fuel utilization and reduced systemic inflammation. Together, these finding suggest MCFAs have the potential to mitigate the development of metabolic disorders and/or facilitate weight loss and maintenance in humans.

## ADDITIONAL INFORMATION SECTION

The authors declare no competing financial interests.

### Author contributions

S.C.S., L.H. and L.Y. designed the research. S.C.S., Q.Q., C.U. and Q.D. performed the research. S.C.S., Q.Q., L.H. and L.Y. analyzed the data. S.C.S., L.H. and L.Y. wrote the article. All authors have read and approved the final version of this manuscript and agree to be accountable for all aspects of the work in ensuring that questions related to the accuracy or integrity of any part of the work are appropriately investigated and resolved. All persons designated as authors qualify for authorship, and all those who qualify for authorship are listed. L.Y. is supported by R01DK108835 and R01DK126817, and S.C.S. and L.H. are supported by R01DE026433.

## FIGURE LEGENDS

**Supplemental Figure 1.**
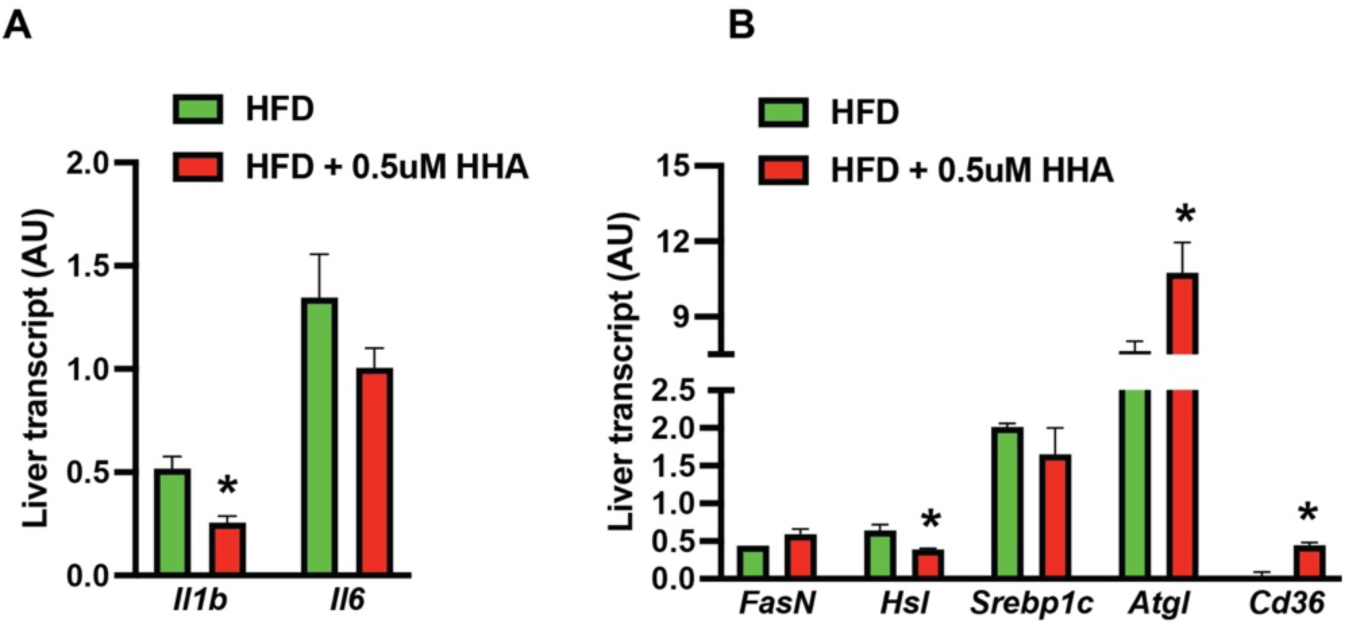
6-HHA modifies HFD-mediated expression of genes in the liver. Mice fed HFD followed by IP injection with control (water) or 6-HHA (0.5 µM) every other day for three weeks. **(A, and B)** RT-qPCR analysis from livers measuring levels of mRNAs encoding the indicated genes. Gene expressions were normalized to *Gapdh*. N = 6 mice/group. All data are presented as means ± SEM. *indicates statistical significance compared to HFD control as determined by multiple Student’s t-test, p<0.05.

## Notes

### Competing Interest Statement

The authors have declared no competing interest.

