## Supplemental Figure 1 for "A Medium Chain Fatty Acid, 6-hydroxyhexanoic acid (6-HHA), Protects Against Obesity and Insulin Resistance"

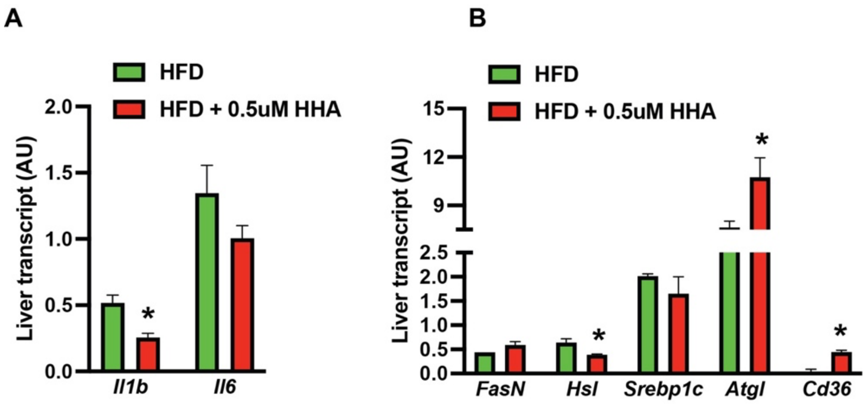


**Supplemental Figure 1. 6-HHA modifies HFD-mediated expression of genes in the liver.** Mice fed HFD followed by IP injection with control (water) or 6-HHA (0.5 µM) every other day for three weeks. **(A, and B)** RT-qPCR analysis from livers measuring levels of mRNAs encoding the indicated genes. Gene expressions were normalized to *Gapdh*. N = 6 mice/group. All data are presented as means ± SEM. *indicates statistical significance compared to HFD control as determined by multiple Student's t-test, $p<0.05$.
